# Boring beetles and super models: mapping the distribution of a new invader

**DOI:** 10.1101/2025.05.15.654388

**Authors:** Andrew Coates, Ben Phillips

## Abstract

The impact of an invasive species is strongly influenced by the extent of its invaded range. Predicting the potential distribution of an invader is thus critical for informing decisions around surveillance, containment, and eradication. Such predictions ideally would come from a mechanistic species distribution model. These models directly describe how aspects of the environment affect organismal fitness, and may also provide insight into population dynamics, such as rates of population growth and spread. Here, we develop a temperature-dependent stage-structured population dynamic model for a globally significant invader, the polyphagous shot-hole borer (PSHB, *Euwallacea fornicatus*). We use this model to predict the beetle’s potential growth rate across its invaded range. Invasive PSHB causes serious damage to urban and agricultural trees, and so the continued expansion of its global distribution is a major concern. We synthesise previous studies to parameterise the model, yielding estimates of population growth as a function of daily temperatures within a host tree. We ran the model over one year using daily climate data across regions with invasive PSHB populations: South Africa, California, Israel and Australia. Our potential distribution maps indicate that PSHB currently occupies the most climatically suitable parts of California and Israel, but is yet to invade the most suitable regions in Australia and South Africa. The model shows the species is capable of very high population growth: in already-invaded locations, the average daily growth rate had an upper bound of ∼ 0.05 (equivalent to population doubling in 14 days). Growth predictions were sensitive to the parameter for adult female mortality during between-host dispersal. This is an uncertain parameter (there is very little data available), but we estimated that for current populations to persist, the loss through dispersal mortality needs to be < 40%. We use our model to predict the maximum spread rate of PSHB (discounting transport by humans) to be 3 km/year. Our model provides insights into the spatial and temporal heterogeneity of invasion risk, which will assist global efforts to manage the ongoing invasion of this species.

## Introduction

Biological invasions are significant drivers of environmental change globally, and come at great costs (Gallardo et al., 2016; Pimentel et al., 2005; Vitousek et al., 1996). Mitigating the impacts of invasive species typically requires direct intervention, such as the containment and/or eradication of populations (Veitch and Clout, 2002). Management actions also carry costs: investment decisions and efficient use of resources are best guided by, among other things, knowledge on the extent of the invasion and how the invasion will progress in the future (Briscoe et al., 2019; Kearney et al., 2008). Because of this, species distribution models (SDMs) have become popular tools for invasive species research and management (Ficetola et al., 2007; Merow et al., 2011; Phillips et al., 2008; Ponti and Gutierrez, 2023). By predicting the potential distribution of an invader across a new landscape, SDMs can be used to generate risk assessments, highlight key regions of concern, and inform decisions on how to deploy control efforts most effectively (Briscoe et al., 2019; Giljohann et al., 2011).

Broadly, there are two approaches to SDMs: correlative and mechanistic (or process-based) models (Briscoe et al., 2019; Buckley et al., 2010; Gallien et al., 2010). The correlative approach is widely used for mapping species distributions, as it is efficient, uses readily-available data, and can be applied across a variety of taxa within a region (Gallien et al., 2010). This method takes occurrence data for a species, correlates it with environmental variables for those sites, and uses this to extrapolate habitat suitability across a landscape according to those environmental predictors.

The inherent assumptions in these models, however, mean that correlative SDMs are error-prone when extrapolating into novel environments and making predictions for non-equilibrium populations – which are defining characteristics of invasion (Briscoe et al., 2019; Elith et al., 2010; Evans et al., 2015; Kearney et al., 2008). Mechanistic SDMs are thus a more suitable alternative for making robust predictions for invasive populations (Phillips et al., 2008; Elith et al., 2010; Gregg et al., 2019; Ponti and Gutierrez, 2023). These models explicitly describe some link between the environment and organismal fitness (e.g., the thermal limits of an organism), and predict distributions by predicting fitness in different environments. An advantage of mechanistic SDMs is that they provide biological explanations behind the expected distributions (Evans et al., 2015; Phillips et al., 2008). In the context of invasions, this allows us a more nuanced understanding of the conditions for population growth and expansion. Here, we apply a mechanistic modelling approach to understanding the potential distribution of a global invader, the polyphagous shothole borer.

The polyphagous shot hole borer (PSHB) has become a global invader in the last two decades. Invasive populations of this wood-boring beetle have established in the United States (California and Hawai’i), Israel, South Africa and Australia (DPIRD, 2024a; Eskalen et al., 2013; Mendel et al., 2012; Paap et al., 2018; Rugman-Jones et al., 2020). Most recently, a population has been detected in Spain (Goldarazena et al., 2025). PSHB belongs to the *Euwallacea fornicatus* species complex (Smith et al., 2019) of ambrosia beetles (Coleoptera: Scolytinae: Xyleborini). Members of the *E. fornicatus* complex are obligate fungus farmers, with PSHB associated primarily with its symbiotic fungus *Fusarium euwallaceae* (Freeman et al., 2013). Adult beetles bore into trees which they inoculate with fungi to use as their sole food source. In their native range in south-east Asia, PSHB predominantly attacks only dead or dying trees (Li et al., 2016). In its introduced range, however, healthy trees are also attacked and suffer severe *Fusarium* dieback as a result of the fungal infection. This has significant ramifications for trees in agriculture, urban environments, and native forests (Coleman et al., 2019; Cook and Broughton, 2023; De Wit et al., 2021; Mendel et al., 2017).

In Australia, for example, the beetle was first detected in 2021, in Perth, Western Australia. Given the beetle’s threat to agriculture and the environment, it has been under active eradication since 2022. The beetle has infected 1000s of trees across more than 100 species (DPIRD, 2024a). Since 2022, more than 3500 PSHB-infested trees have been cut down in Perth in an attempt to control the beetle as part of the $AUD40m eradication response (Burmas, 2024), and a quarantine area has been established around the current range (DPIRD, 2024a) to help contain the beetle’s spread.

### Modelling PSHB

Two recent studies have constructed SDMs for PSHB in Australia, to assist with eradication efforts (Li et al., 2024; Warnakula and Parsons, 2024). These efforts both used a correlative model approach, in which known PSHB occurrences were correlated with climatic variables (temperature and precipitation data) and used to extrapolate the suitability of habitats in Australia. Given the uncertainties associated with using correlative SDMs in an invasion setting (Briscoe et al., 2019), supplementing these predictions with a mechanistic approach seems warranted. Using a range of methods will improve confidence in which areas have a high risk of invasion, and where further research is required.

A drawback to mechanistic SDMs is that model development is more demanding: these models typically require physiological data specific to the focal species to delineate aspects of the fundamental niche. Fortunately, several studies have already gathered useful data on the thermal responses of the *E. wallacea* species complex (Dodge, 2019; Umeda and Paine, 2019; Walgama and Zalucki, 2007).

This study builds a temperature-dependent stage-structured population model for PSHB, as well as a simple sub-model that predicts tree temperature (the microhabitat the beetle inhabits). The result is a model that, for a given location, estimates daily population growth accounting for the microhabitat and full thermal performance curve of the organism. Such a model can be applied across space to estimate the species distribution, but also provides a richer understanding of the organism and its limits. We apply the model to PSHB’s non-native range: Australia, South Africa, California and Israel. We focus on Australia in particular as this is the most recent invasion, and the one currently undergoing intensive control. In all these areas, the model directly estimates population growth rate, which is used to predict the distribution but can also be used (in conjunction with dispersal estimates) to estimate the spread rate of the invasion. The model’s daily time steps also allow us to examine seasonal variation in growth, which provides insight into the conditions limiting the distribution, but also provides managers with clear guidance about when surveillance will be most effective. Finally, by cross referencing the model against locations where the beetle is known to persist, we can learn about unmeasured model parameters, which provide insight into population control.

## Methods

We constructed a discrete time, stage-structured matrix model for a PSHB population. The model tracked the number of female PSHB through time (in days, *t*) occupying a host tree. We modelled only females, as they are the drivers of invasion for several reasons. First, the PSHB sex ratio is heavily female-skewed, ranging 3:1 to 14:1 (Cooperband et al., 2016; Gadd, 1941). Second, there is little mate limitation, as females mate with their siblings or, in the absence of siblings, can produce male offspring asexually to mate with (Cooperband et al., 2016). Third, it is females that disperse and establish new colonies; males usually remain in their natal gallery for their entire life (Cooperband et al., 2016).

We partitioned the female population into three stages, based on life history: the ‘juvenile’, ‘preoviposition’ and ‘adult’ stages; *i* = {*J, P, A*}. The juvenile stage, *J*, encompasses the egg, larva and pupa phases. The preoviposition stage, *P*, represents new adults as they emerge from their natal gallery and establish a new colony, either on the same host tree or, after dispersing, on a new host. After the preoviposition period, females then become adults, *A*, which produce new offspring. The number of individuals in each stage is calculated for each time step, *t*, according to these transition equations:

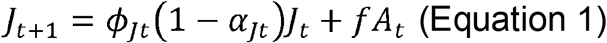

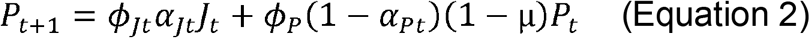

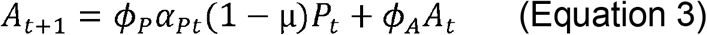

The matrix equation parameters are summarised in Table 1. The transition probabilities for each stage, *α*_*it*_, are functions of daily temperature and describe the daily proportion of individuals that transition from one stage to the next. The parameters Φ_*i*_ are the daily survival probabilities for each stage. Juvenile survival, Φ_*Jt*_, is a function of daily temperature. Other survival probabilities are set to be constant (independent of temperature), according to a background mortality rate. The proportion of pre-ovipositing adults, *P*, lost to mortality during dispersal to a new host is given by *µ*. The parameter *f* is the mean daily fecundity (*i*.*e*., number of female offspring produced by an adult female per day).

**Table 1.** Description and values of indices and life history parameters used in the matrix population model (Equations 1 – 4).

| Indices | Description | Value |
| --- | --- | --- |
| $t$ | Time-step | Represents 1 day. |
| $i$ | Life stage | Juvenile, Pre-oviposition or Adult:<br>$i = \{J, P, A\}$ |
| **Life history parameters** |  |  |
| $\alpha$ | Transition probability (to next life stage) | Temperature-dependent |
| $\phi$ | Survival probability | $\phi_J$ = temperature dependent<br>$\phi_P = 0.97$<br>$\phi_A = 0.97$ |
| $f$ | Fecundity (eggs per adult per day) | $f = 0.69$ |
| $\mu$ | Probability of $P$ mortality through dispersal | $\mu = 0 - 0.95$ (ranged across simulations) |

Equations 1 – 3 are captured within a matrix recursion equation:

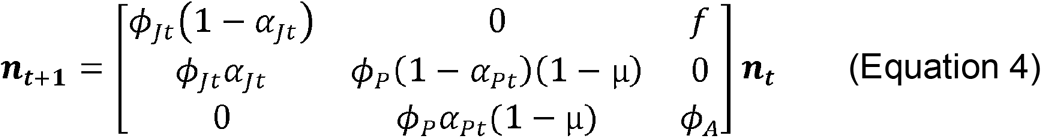

Which calculates through time the number of PSHB in each stage, given by the column vector ***n*** = {*j, P, A*}.

### Estimating temperature-dependent transition probabilities, α

The temperature-dependent development rates of the *E. wallacea* species complex have been demonstrated in several laboratory studies (Dodge, 2019; Umeda and Paine, 2019; Walgama and Zalucki, 2007). Successful development of *E. wallacea* occurs within a thermal window approximately 13–33 °C, with an optimum developmental temperature of approximately 28° (Dodge, 2019).

To generate functions for temperature-dependent transition probabilities (α), we fit temperature performance curves (TPCs) to empirical PSHB data. TPCs describe the relationship between temperature, *T*, and performance, *P*, which in this instance is the probability of transition. As in Phillips et al., (2014), we describe a TPC function that is a meeting of two Gaussian functions:

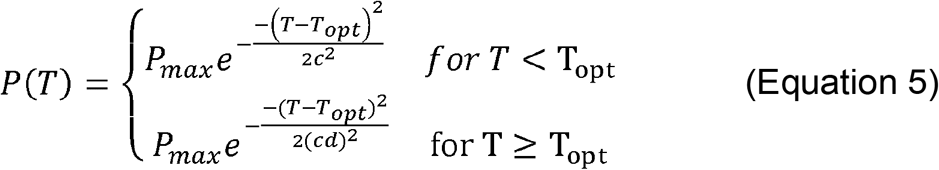

Where *P*_*max*_ is the maximum performance value and *T*_*opt*_ is the optimum temperature (*i*.*e*., the temperature at *P*_*max*_). Performance increases with temperature up to *T*_*opt*_, according to *c*. At temperatures above *T*_*opt*_, performance decreases, as given by *d*.

TPCs for α_*J*_ and α_*P*_ were fitted to empirical data on PSHB development rates at different temperatures (from Umeda & Paine, 2019; and Walgama & Zalucki, 2007). The fitting of these curves and the parameter estimates for Equation 5 are described in detail in the Supplementary Material. When running the population model, tree temperature was input as *T* into the parameter-specific TPC functions (Equation 5) and used to calculate the values of α_*J*_ and α_*P*_ for that time-step.

### Estimating survival probabilities, □

Juvenile survival, □_*J*_, is also temperature-dependent (Walgama and Zalucki, 2007). We calculated daily survival probabilities at different temperatures from the empirical data given in Walgama and Zalucki, (2007). As with α, a function for temperature-dependent □_*J*_ was generated by fitting a TPC to these data (as described above, Supplementary Material). The mortality rates of adult stages, □_*P*_ and □_*A*_, appear to be less affected by temperature, and so we assumed them to be constant. Gadd (1941) estimated the mean laying lifespan of an adult females as 32 days (maximum 49 days). This yields a per day mortality rate of 1/32, which equates to a daily survival probability, □_*A*_, of 0.97. In the absence of any other data, we assume □_*P*_ = □_*A*_.

### Estimating fecundity, f

Gadd (1941) estimated the rate of egg laying at 0.927 eggs per day per female. In this study, 75% of offspring were female. Hence, the daily production of female eggs per adult female was parameterised as *f* = 0.69. In nature, this is almost certainly also temperature dependent, but in the absence of data on laying rate as a function of temperature, we keep this value constant in the model.

### Estimating dispersal mortality, μ

A proportion of individuals in the preoviposition stage, are expected to be lost from the system as a result of dispersal to a new host, given by the parameter *µ*. This parameter captures the net loss of females through emigration, as well as mortality during dispersal, which reduces the immigration of females from other trees. There is little information in the literature for us to estimate *µ*. To begin, we run the simulations with *µ* = 0 (i.e., assuming negligible mortality from dispersal). We then ran suites of simulations with *µ* ranging 0 – 0.95 for select invaded locations, to refine the potential range of this parameter (described below). Based on these simulations, we also generated distribution maps with *µ* = 0.2 and 0.4.

### Distribution maps

To assess the spatial distribution of PSHB suitability across Australia, we used daily weather data obtained from the Queensland Government SILO climate database (https://www.longpaddock.qld.gov.au/silo/; accessed via WeatherOz: Pires et al., 2024), which provides an extensive dataset of interpolated meteorological variables for all of Australia. We ran the population model independently across a grid of coordinates covering all of mainland Australia and Tasmania, with a resolution of 0.1 × 0.1 degrees (equivalent to ∼ 10 × 10 km; for a total of 139,090 locations). For each coordinate, daily temperature and humidity data were taken for 2013 – 2023, and averaged values across the 10 years to generate mean daily values for every day of the year.

For other invaded countries – South Africa, USA (California) and Israel – we used climate data obtained from NASA’s POWER (Prediction Of Worldwide Energy Resources) global climate database (Sparks, 2018). As above, daily temperature and humidity values were averaged across a 10-year period (2013 – 2023), for coordinates encompassing a 0.1° x 0.1° grid over these locations. The spatial resolution of the POWER database is coarser than that for SILO (0.5° latitude x 0.625° longitude).

For each coordinate in the grid the population model was run for one year (*t* = 1 – 366, beginning January 1^st^). We started the simulations with a founding population of one adult female. Outputs included the number of individuals each day and the population growth rate per life stage per day. To produce distribution maps, we calculated for each coordinate the mean daily population growth rate for the adult life stage over the whole year, 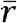:

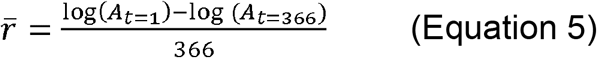

We also examined seasonal variation in population dynamics by looking at daily growth rates at key locations; typically major cities within the region of interest. For Australia, for example, we focused on daily rates in 10 cities across the country (the capital cities, plus Cairns and Broome; **Figure 5**) to look at seasonal variation in suitability. The exact coordinates used were for the cities’ Botanic Gardens (or the golf club, Broome); central urban locations expected to have an abundance of potential host trees.

### Tree temperature predictions

The PSHB life cycle takes place almost entirely inside the host tree, within galleries excavated into the xylem tissue. The microhabitat experienced by wood-boring beetles inside trees is often several degrees warmer than the ambient air temperature (Keena and Moore, 2010; Logan and Powell, 2001; Trần et al., 2007). The wood can also buffer against extreme temperature changes (Formby et al., 2018; Powell, 1967). To describe this microenvironment, we created a function that predicted daily tree temperature (*T*_*tree*_) as a weighted function of the maximum air temperature (*T*_*max*_) and the temperature of the soil 1 metre underground (*T*_*soil*_), according to:

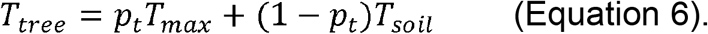

This assumes that the sapwood temperature always falls between the maximum air temperature and the soil temperature. Soil temperature determines the temperature of the water drawn up through the roots, which in turn influences wood temperature (Powell, 1967). The relative effect of soil temperature increases with transpiration rate (due to a larger volume of water being drawn into the xylem), and transpiration rate is inversely proportional to ambient humidity (Jones, 2014). Therefore, in our model, the relative weight of air and soil at determining *T*_*tree*_ is given by *p*_*t*_, which can take a value between 0 and 1, and is a function of the relative humidity, *H*. Specifically, we assumed a linear relationship (on the logit scale) with humidity:

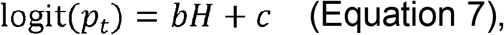

where *b* and *c* are parameters of a linear model. We fit the model to a data set for King’s Park, Perth, that combined daily air temperature and humidity data (from SILO, as above) with tree temperature data recorded using sap flow loggers (data provided by E. Veneklaas; W. Lewandrowski). The sap flow data sets included half-hourly measurements from temperature probes, taken over a total of 15 months (converted to average temperature per day of the year). We calculated soil temperature as the rolling average of the mean air temperature over the last 30 days. We found this to be a good approximation, when tested against a more complex soil microclimate model (Kearney and Porter, 2019). We fitted the tree temperature function to the climate and sap flow data in a Bayesian inference framework, using HMC implemented in the greta package (Golding, 2019). This process of estimating parameters *b* and *c* is described in detail in the Supplementary Material.

### Testing the model with already invaded locations

To assess whether our model was producing reasonable predictions, we examined outputs for a set of locations where invasive PSHB have established populations. We selected four locations in South Africa from the database of PSHB observations at FABI, (2025): in Johannesburg, Durban, George and Cape Town. These were urban records, chosen to represent four main clusters of PSHB observations in South Africa (Van Rooyen et al., 2021). Records of PSHB in California were obtained from ANR, (2025). We selected three locations – Laguna Beach, Santa Paula, San Marino – covering three different counties – Orange County, Ventura County, LA County, respectively. Laguna Beach represented the southernmost extent of the PSHB distribution; Santa Paula was the northernmost extent. San Marino, which had many records of the pest (ANR, 2025), was chosen to represent the middle of the range. PSHB records for specific sites in Israel were not available, although we did examine outputs for Tel-Aviv, as Freeman et al., (2013) collected *Fusarium* fungus from several areas surrounding the city.

For the invaded coordinates, we ran the model under different *µ* values, ranging 0 – 0.95 (at 0.05 intervals). We then looked at the maximum *µ* values under which a PSHB population would be predicted to persist (*i*.*e*., where 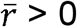).

Simulations were run on R version 4.3.0 using the Pawsey Setonix supercomputer (Pawsey Supercomputing Research Centre, 2023).

### Estimating invasion speed

It is possible to estimate an invasion’s spread rate according to 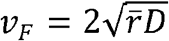, where *v*_*F*_ is the invasion speed (the Fisher velocity), 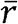 is the population growth rate and *D* is a diffusion coefficient (Bîrzu et al., 2018; Fisher, 1937). The Fisher velocity is estimated under the assumptions of continuous space and time, a constant gaussian dispersal kernel, and logistic (or exponential) population growth. Many expected breaches of these assumptions (*e*.*g*., a discontinuous space; Allee effects) would result in an actual velocity lower than *v*_*F*_ (Phillips, 2025). On the other hand, a dispersal kernel with long tails would cause a velocity higher than *v*_*F*_ (Kot et al., 1996). Nonetheless, choosing the highest value of 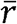 produced by our model (*i*.*e*., population growth where *µ* = 0) allows us to determine a likely upper bound on PSHB spread rate at a location (assuming a gaussian kernel). To estimate a value for the diffusion coefficient, *D*, we used data from an *E. fornicatus* mark-release-recapture experiment (Owens et al., 2019). This dataset (pers. comm. D. Owens and P. Kendra,) describes the number of adult females recaptured by traps arranged 15 – 130 m from a release point. We fit this dispersal data to a 2D Gaussian distribution using, again, the greta Bayesian inference machinery (Golding, 2019) to estimate the mean squared displacement, *σ* (details in Supplementary Material). Given = 25.7 and *t* = 1 (1-day increments), the daily diffusion coefficient was calculated as 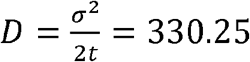 (Phillips, 2025; Shigesada and Kawasaki, 1997).

## Results

### Testing model for invaded locations

Our model predicted relatively high suitability for locations known to have invasive PSHB populations (in South Africa, California and Israel). When *µ* was set to zero, daily population growth was positive for most of the year at all locations (Figures 1 – 3). Mean growth rates ranged from = 0.037 (Cape Town) to = 0.057 (Durban). Unsurprisingly, population growth decreased as the parameter value for *µ* (dispersal mortality) was increased (Figure 4). The predicted upper limits for *µ* (the maximum values under which was positive) at invaded locations was 0.4 – 0.5 (Figure 4). This suggests that the average daily loss of dispersers must be < 40% for PSHB populations to be maintained.

**Figure 1.**
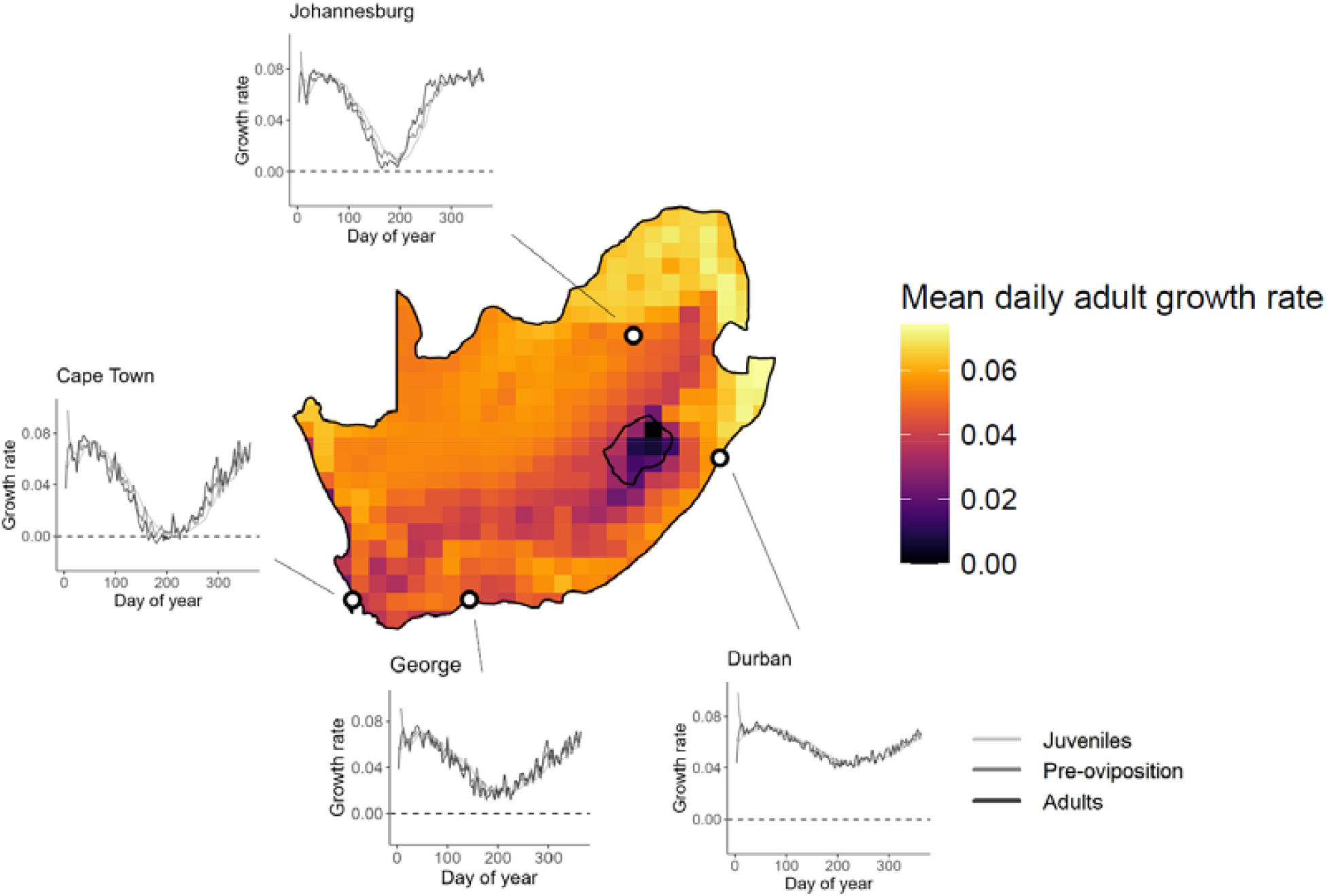
PSHB climate suitability distribution map for South Africa and Lesotho (given as mean daily adult population growth rate). Also given are the daily population growth rates over one year for four locations within the current PSHB distribution. Predictions are for *µ* = 0.

Potential distribution maps for South Africa, California and Israel are given in Figures 1 –3. In South Africa, the highest suitability occurred in the north-east of the country (Figure 1). The distinctive region of low suitability around Lesotho corresponds to areas of high elevation (Moore et al., 2009). The current PSHB range in California (ANR, 2025) is situated in the region with the highest predicted suitability (Figure 2). Suitability was lowest in Yosemite National Park. Conditions were highly favourable across Israel (values ranged 0.031 – 0.057), especially along the coastline (Figure 3). A seasonal cycle of population growth was observed in all invaded locations, with the highest growth occurring in summer. The exception to this was in Tel-Aviv, Israel, where population growth slowed in the middle of summer, when mean tree temperatures exceeded 30° C.

**Figure 2.**
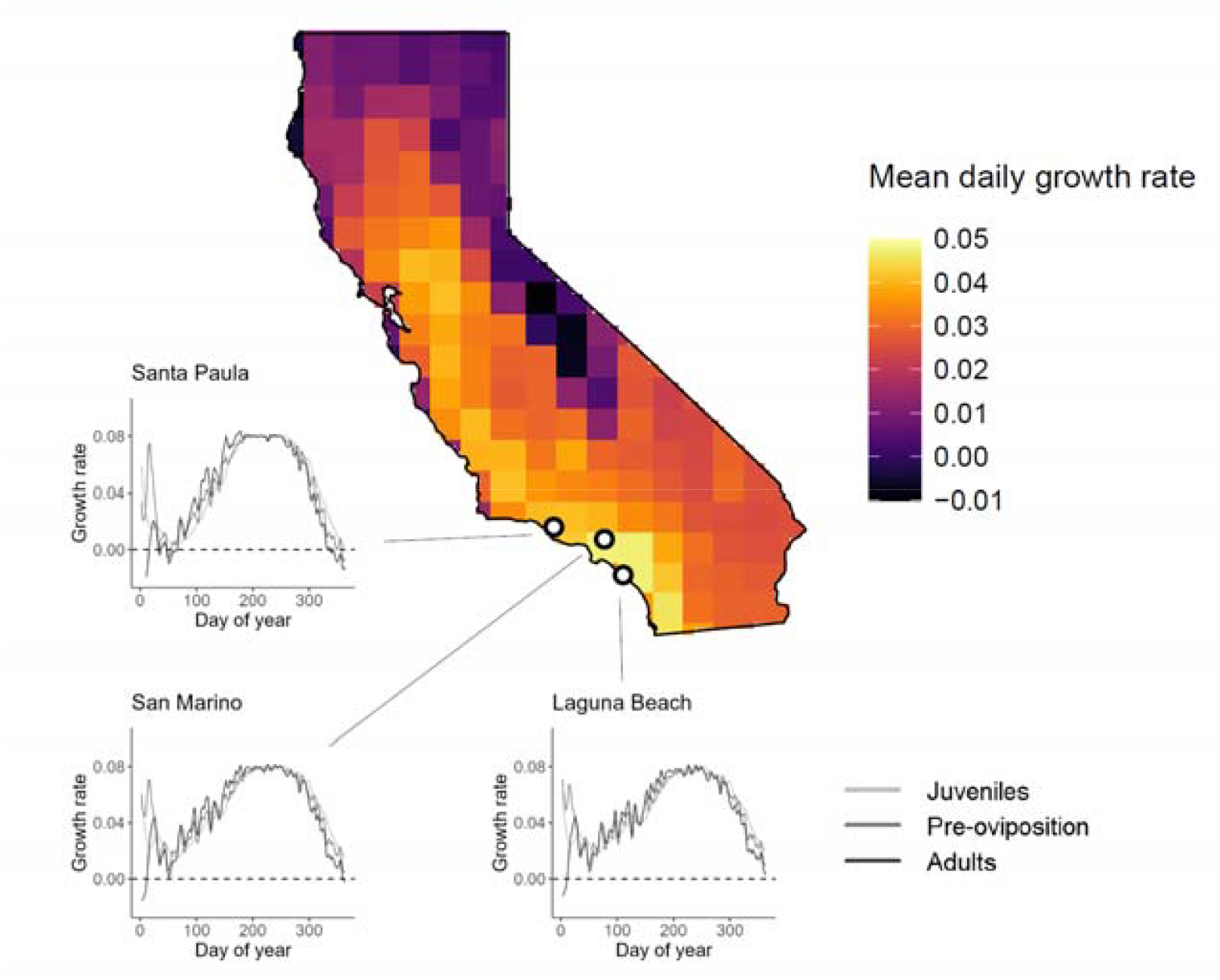
PSHB climate suitability distribution map for California, USA (given as mean daily adult population growth rate). Also given are the daily population growth rates over one year three locations within the current PSHB distribution. Predictions are for *µ* = 0.

**Figure 3.**
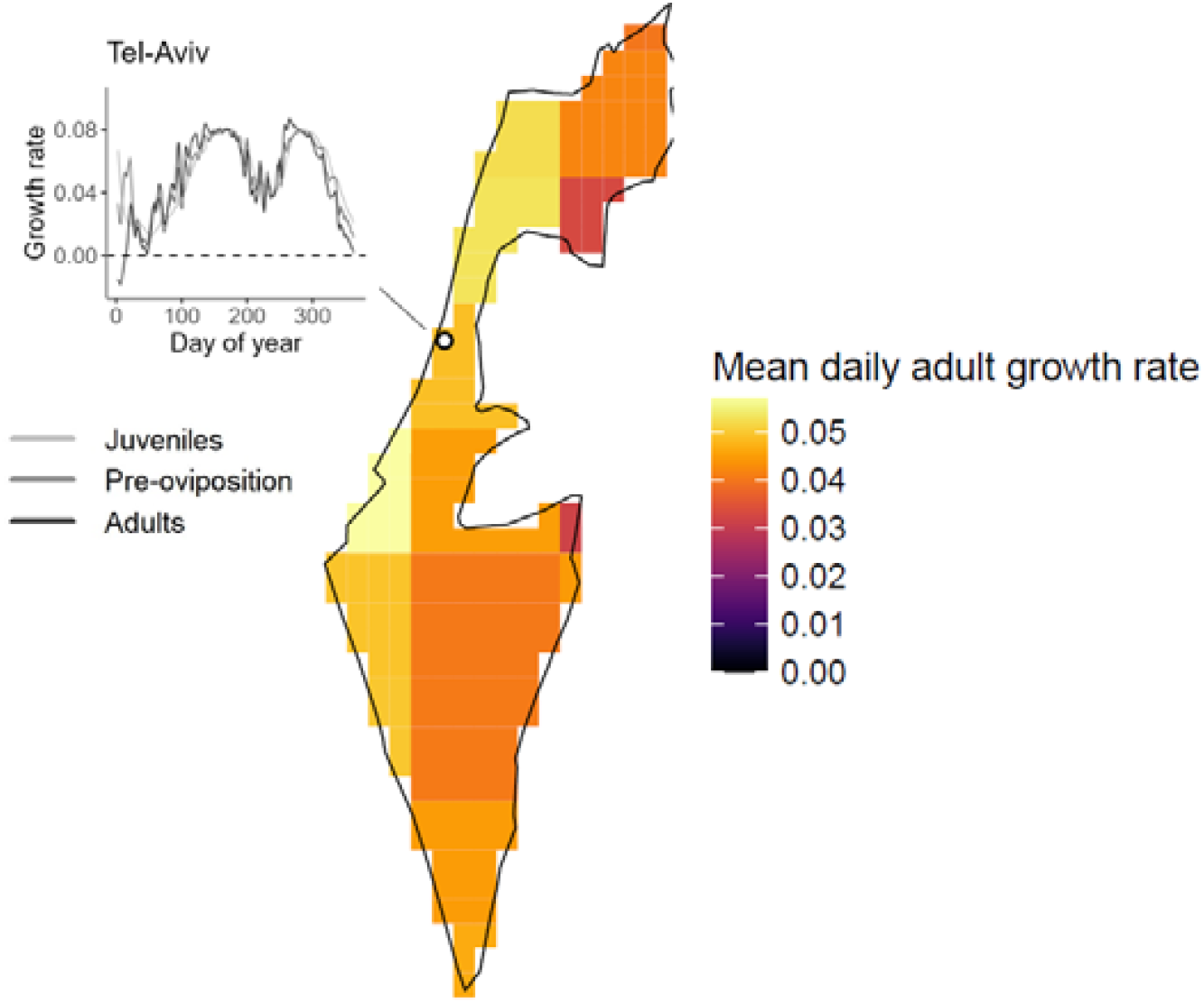
PSHB climate suitability distribution map for Israel (given as mean daily adult population growth rate). Also given are the daily population growth rates over one year for Tel-Aviv. Predictions are for *µ* = 0.

**Figure 4.**
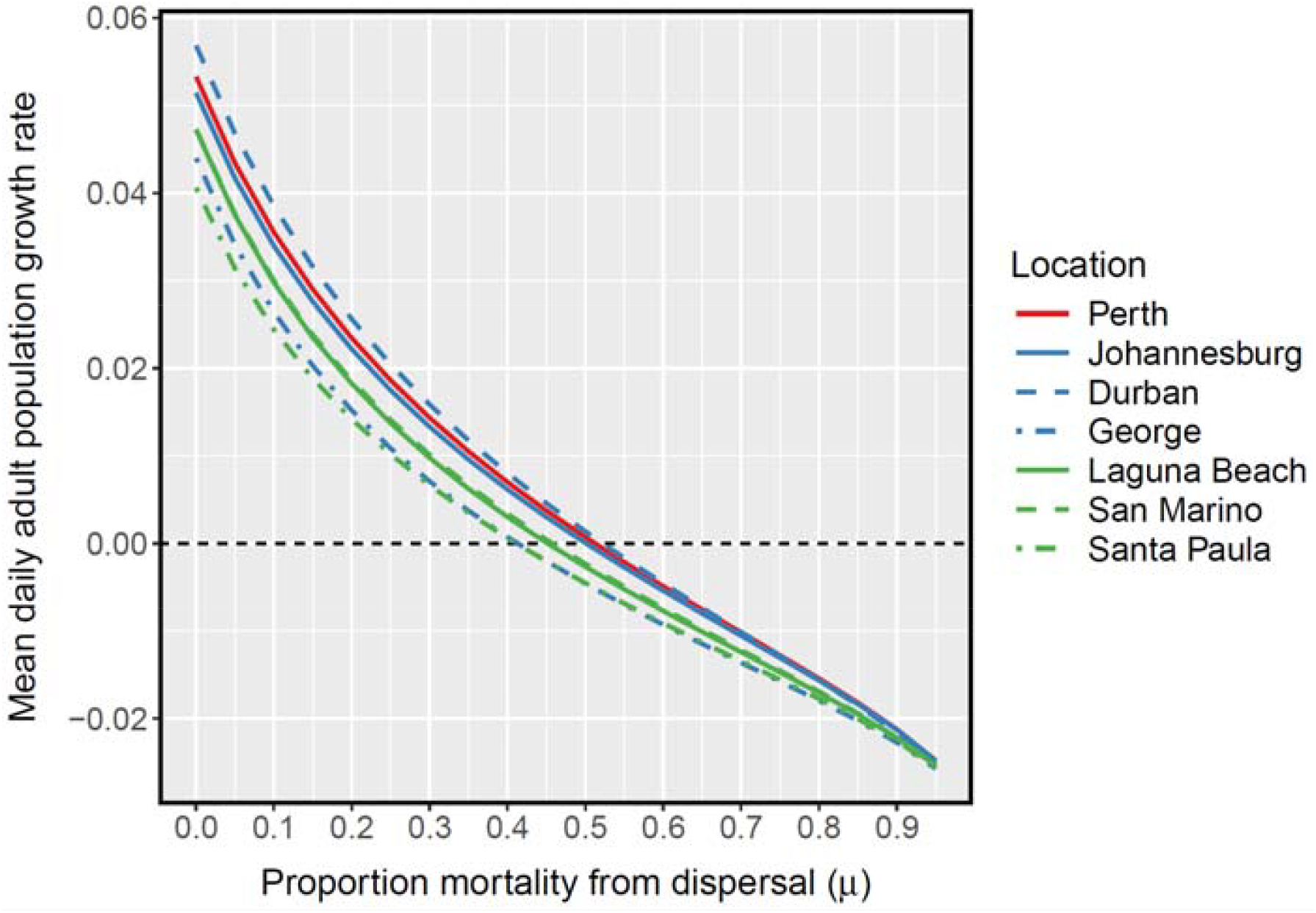
Mean daily population growth rate (of adults over one years) as a function of the parameter for dispersal mortality, *µ* (ranging from 0 – 0.95). Models run using climate data for locations with known PSHB populations (in Australia, red; South Africa, blue; California, green).

### Potential distribution in Australia

The potential PSHB distribution maps for Australia are given in Figure 5 –**Figure 6**. The current PSHB distribution in Perth, WA, is in a region of relatively high suitability, with a mean population growth rate of 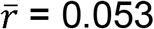 (assuming no dispersal mortality, *µ* = 0). This region of high population growth extended north from Perth along the coastline (up to Shark Bay). The high-suitability climate also extended south-east into the Nullarbor plain (southern Western Australia and South Australia). Much of the east coast had favourable conditions: the population dynamics predicted for Sydney were very similar to that of Perth, and the suitability increased further northwards. The north-eastern coastline of Australia was the most favourable environment for PSHB, with growth rates of 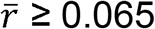 from Brisbane to Cairns and into Cape York (Figure 5).

**Figure 5.**
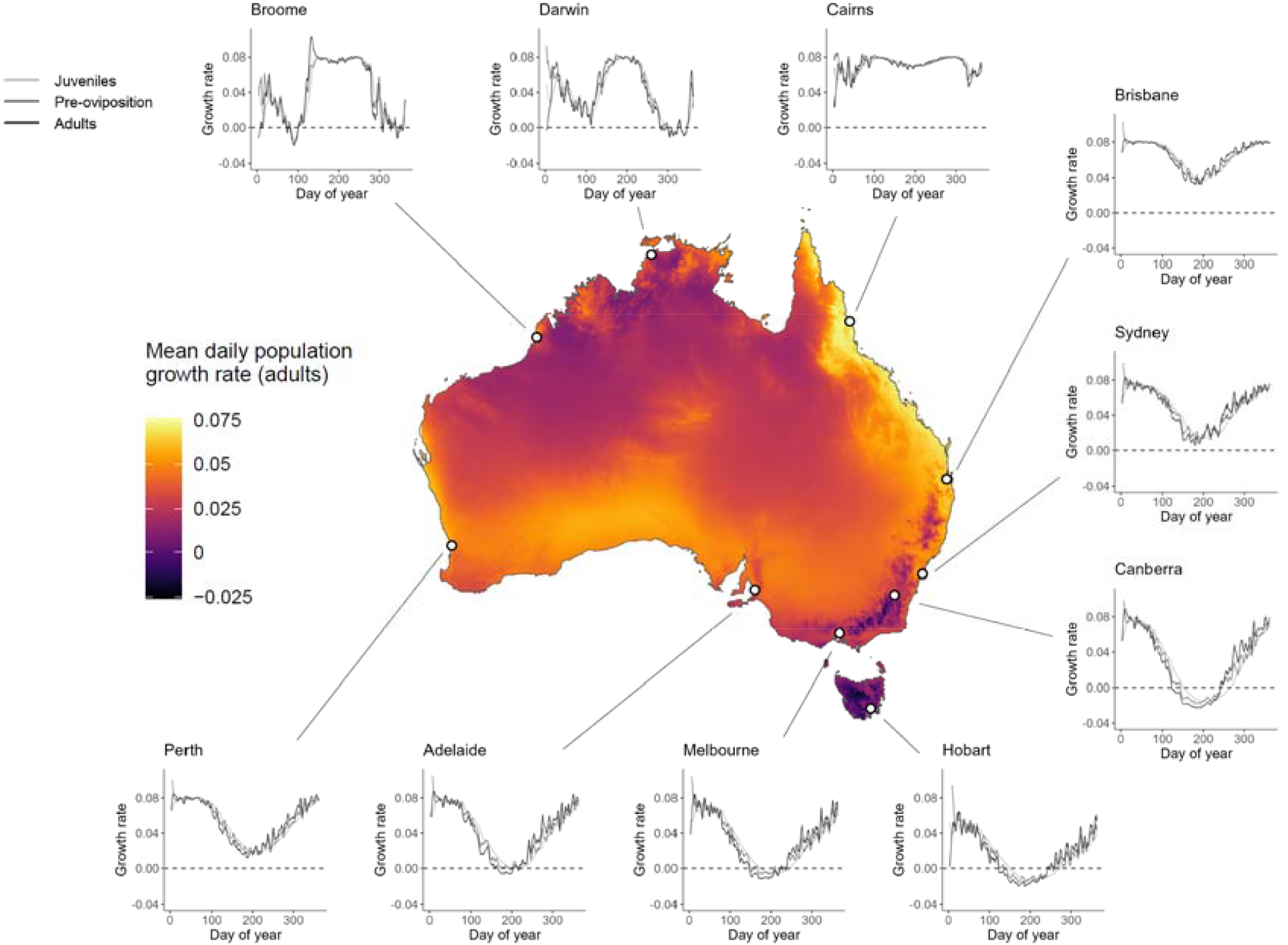
Predicted habitat suitability for PSHB in Australia, according to the mean daily population growth rate (of adult females) over one year. The growth rate shown is predicted by the mechanistic model with no loss of pre-adults through dispersal (µ = 0). The most suitable locations have highest average population growth rates (yellow). Map resolution of 0.1 × 0.1 degrees. Also includes daily growth rates (juveniles, pre-oviposition adults, adults; light grey, dark grey, black) over one year for major Australian population centres.

**Figure 6.**
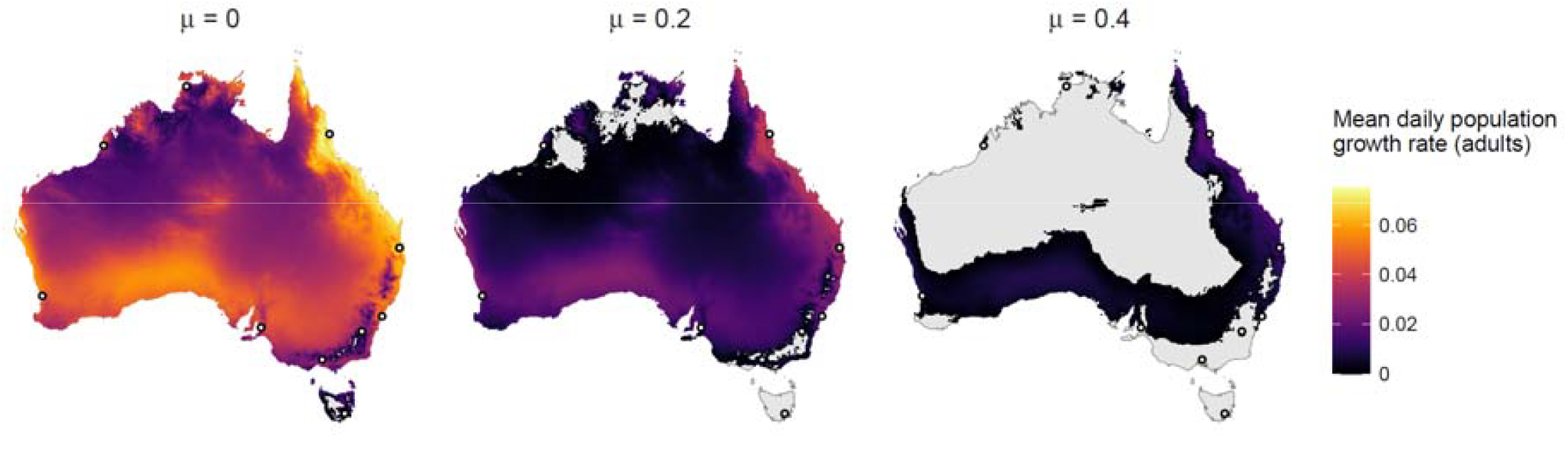
Predicted habitat suitability for PSHB in Australia, according to the mean daily population growth rate (of adult females) over one year. Predicted distributions shown under different assumptions of dispersal mortality: *µ* = 0, 0.2 and 0.4. Regions with negative mean growth rates (where populations are not expected to persist) are uncoloured.

The least suitable regions were in the south-east of the continent: in Tasmania and the Victoria/NSW Alpine region. These were the two areas where mean population growth, 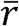, was predicted to be below zero, even without dispersal mortality (*µ* = 0). The potential distribution of PSHB (*i*.*e*., where predicted population growth was positive) contracted considerably as the value of *µ* was increased (Figure 6). At *µ* = 0.4, the majority of Australia was predicted unsuitable for PSHB invasion. However, there is a continuous band of favourable climate linking the west and east of the continent.

In most Australian cities examined, there was peak population growth in the summer (**Figure 5**). Over winter growth rates decreased, and in the southernmost cities populations declined (i.e., negative population growth). The exceptions to this were cities in the tropics (Darwin, Broome and Cairns) which have a characteristic wet-dry seasonality. In these locations, population growth tended to be highest during the cooler dry season (the austral winter). During the warmer wet season, when temperatures exceeded 30 °C, population growth was lower (and occasionally negative).

### Estimating spread rate

We solved the equation for the Fisher velocity, 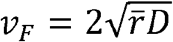, to estimate the speed of the PSHB invasion in Perth, given 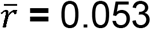 (the mean daily population growth rate) and *D* = 330.25 (from Owens et al., 2019; Supp. Material). This gives *v*_*F*_ = 8.4 m/day, equivalent to 3.1 km/year. At 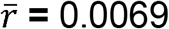 (mean population growth at King’s Park under *µ* = 0.4), the estimated velocity was 1.1 km/year.

## Discussion

In this paper, we use a temperature-dependent stage-structured population model to integrate environmental temperatures over a year into an estimate of organismal fitness. We apply this approach to a global invader — the polyphagous shothole borer (PSHB) — and use it to estimate the species’ potential distribution in its non-native range. Our model also provides valuable insight into the temporal and spatial dynamics of the invasion.

### PSHB distributions

In Australia, the polyphagous shothole borer (PSHB) is currently restricted to metropolitan Perth, WA, but there are concerns that it will continue to expand its range and inflict substantial damage to trees over a large scale (DPIRD, 2024a). Our species distribution model suggests these concerns about range expansion are well founded. Our model predicts that large swathes of Australia have the thermal conditions suitable for PSHB to establish in. The west coast of WA (including Perth) was one of the most favourable regions in all of Australia (**Figure 5**), which likely facilitated rapid PSHB establishment following introduction. We expect the population to advance most quickly where population growth is highest, as predicted by the Fisher-KPP equation (Phillips, 2025). Most notably, the coastline north of Perth is likely a key invasion corridor. Where population sizes are amplified, we also expect the impact on host communities to increase. As such, the regions of high PSHB suitability predicted by our model would be good candidates for prioritising control efforts. Beyond Perth, our maps identified other regions with high suitability for PSHB establishment – most notably the east coast of Australia. As an agricultural pest, PSHB is most associated with avocado orchards (Mendel et al., 2012). In Australia, the majority of avocado production occurs in Queensland and WA, in the regions predicted to have high PSHB favourability (Avocados Australia, 2024). Given the potential for the long-distance, human-mediated transport, strict biosecurity measures will be necessary to prevent secondary introductions of PSHB into new parts of Australia.

The current PSHB range in California (ANR, 2025) is situated in the region with the highest thermal suitability, as predicted by the model. Conditions appear most favourable for the invasion to extend further south, where there is already an established population of the closely-related Kuroshio shot hole borer (KSHB, within the same *E. wallacea* complex). By contrast, the highest-suitability regions in South Africa currently have few records of PSHB (FABI, 2025), although this may speak to a lower monitoring effort in those areas. The coarse spatial resolution of the climate data meant that we could not distinguish fine spatial differences in Israel, although generally the climate was most suitable along the coast.

### Spread rate

In the simplest model of an invasion, we expect invasions to reach an equilibrium where the range expands at a constant speed, given by the Fisher velocity, *v*_*F*_ (Phillips, 2025). We estimated that the PSHB invasion could advance at a rate up to ∼3.1 km per year, which is relatively slow compared to some invading species (*e*.*g*., the cane toad range edge in Australia expands at a rate of 50km per year; Urban et al., 2008). However, *v*_*F*_ was estimated under multiple assumptions, including continuous space and time, a constant gaussian dispersal kernel (no human-mediated long-distance dispersal), and logistic population growth. Breaches of some of these assumptions (*e*.*g*., a discontinuous space; Allee effects) are expected and would result in slower invasion speed (Phillips, 2025). Conversely, a long-tailed dispersal kernel (including wind-mediated or human-mediated long-distance transport) is possible for this beetle (Owens et al., 2019) which can result in the invasion speeds well beyond *v*_*F*_ (Kot et al., 1996). We estimated the diffusion coefficient using field trial data (Owens et al., 2019) as this is most representative of natural dispersal patterns, but longer flight distances have been recorded using a flight mill (Owens et al., 2019). The beetle’s presence on Rottnest Island, WA, which is > 30 km off the mainland, may also speak to long-distance dispersal of adults on prevailing winds (alternatively, to human transportation). Temperature influences walking performance of PSHB (Pienaar et al., 2025), and this may extend to temperature-dependent flight distance. The estimated spread rate of 3 km per year did, however, assume population growth without dispersal mortality (*µ* = 0), and so is likely an upper bound as far as population growth is concerned. With the more conservative estimate of 40% dispersal mortality, unassisted invasion speed was estimated to be much slower, at 1.1 km per year.

The long-distance expansion of the PSHB distribution is likely to occur through human-mediated dispersal, via the transport of infested timber. This is the probable explanation for the current confirmed distribution in South Africa, where populations are separated by hundreds of kilometres (Van Rooyen et al., 2021). Our results reinforce the importance of the containment measures that are in place around metropolitan Perth, to prevent secondary introductions to new locations (DPIRD, 2024a).

### Comparing SDMs

Comparing and contrasting outputs from different SDMs can build a clearer picture of where there is higher certainty surrounding predictions, and where further assessment is needed to determine invasion risk. In the future, outputs from different methods can be synthesised to enhance predictive power. For example, correlative models can be improved by integrating findings from mechanistic models in hybrid distribution models (Gallien et al., 2010; Kumar et al., 2014; Rodríguez et al., 2019).

Our predictions were broadly consistent with previous modelled distributions of PSHB across Australia. Figure 7b is the distribution from a correlative SDM using the GAM method (Li et al., 2024), recreated from the raster file provided by X. Li. Another distribution produced using the correlative Climatch analysis can be found at Warnakula and Parsons, (2024). The potential distribution of PSHB in Australia has also been predicted by Umeda, (2017), using the CLIMEX program, which incorporates physiological processes (in this instance, temperature-dependent development rates) into a correlative SDM framework. This map has been recreated for Figure 7c.

**Figure 7.**
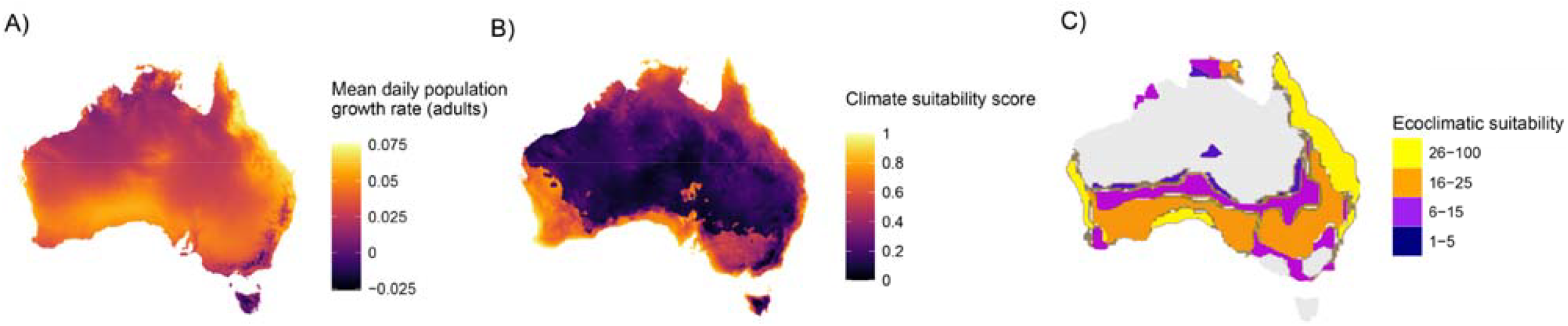
Relative suitability maps for PSHB in Australia derived from A) the mechanistic approach in this study; B) a GAM correlative SDM (from Li et al., 2024); C) CLIMEX model (from Umeda, 2017).

All approaches highlighted regions of very high suitability along the east and west coasts. Our predictions of high suitability on the southern coast (along the Great Australian Bight) were similar to the GAM output (Figure 7), although spatial patterns in other areas were more similar to the Climatch analysis (Warnakula and Parsons, 2024). With the mechanistic component of CLIMEX added, the suitability map in Umeda (2017) had many similarities to those produced by our model – most notably the continuous band of higher-suitability habitat in Australia, running from the west coast, across the south, and up to northern Queensland (Fig. 2C; Fig. 7C). Our distribution map for California was also similar to previous model predictions (Lynch et al., 2025; Umeda, 2017), with high suitability in the south of the state and through the Central Valley.

An advantage of mechanistic models is the ability to explore underlying factors that determine the potential distribution. In Australia, the limiting factor to PSHB invasion in the south was extended periods of low temperature. During winter in the coldest areas (Tasmania and the Victoria/NSW alpine region), populations declined as beetles were lost through mortality faster than could be replenished through maturation and reproduction. The distributions predicted for South Africa and California were also restricted primarily by low temperatures (see Supp. Fig. 4), especially in high-altitude regions. In northern Australia, by contrast, high temperatures were the main hurdle to PSHB establishment. In much of central and northern Australia, daily tree temperatures exceeded 30 °C for extended periods, which increased juvenile mortality and stymied juvenile development. Temperatures above the thermal optima for growth and survival similarly reduced suitability in the south-eastern corner of California (in the Colorado and Mojave deserts). This differed from the results of a degree-day model (that didn’t take survival rates into account), where higher temperatures were directly proportional to higher suitability.(Lynch et al., 2025; Umeda, 2017).

The strength of mechanistic predictions increases as more life-history axes are added to the model (Abarca et al., 2024). By incorporating two separate, temperature-dependent processes (development rate and survival) to our model, our outputs were likely better reflections of reality compared to modelling just one. Another advantage of our model is that distributions are generated directly from a population model, which provides quantitative measures of habitat suitability (such as population growth rate at daily and yearly scales). These outputs can also be used to predict spread rates and other demographic processes. This is in contrast to correlative models, which produce suitability indices that are more difficult to define and to use to understand ranges (Buckley et al., 2010).

Our approach also allows for temporal variation in suitability to be assessed, which is useful for a number of reasons. How PSHB numbers in an affected area increase or decline with the seasons – and the resulting impact on tree communities – can be predicted, as can how the PSHB range expands or contracts through time. It is also valuable information for monitoring. PSHB monitoring is conducted mainly using baited traps (DPIRD, 2024a; Kendra et al., 2020), but this only targets adult females during dispersal. The chances of successfully detecting PSHB occurrence will be higher if traps are set during periods predicted to have high growth rates of emerging females. In Perth, for example, our model suggests that detection rates will be highest at the end of summer, following an extended period of high population growth. Detection efforts over winter, by contrast, are likely to be substantially less effective.

### Limitations of the model

There are many abiotic factors that shape the fundamental niche of a species, and focusing only on temperature is likely to overestimate the potential distribution (Buckley et al., 2010; Kearney et al., 2008). The influence of other abiotic factors may be limited for PSHB, however, given the sheltered environment of the tree galleries they live in (the exception being for dispersing adults where dispersal rates are thought to be lower during periods of rainfall; Liu et al., 2022). Biotic factors were also absent from our model, the most crucial being the availability of host trees. For example, the actual suitability of the Nullarbor Plain is likely to be less than that predicted by our model – although the thermal environment supports PSHB, the region is famous for its low tree density. There is a growing list of the plant species in Australia known to be hosts for PSHB (DPIRD, 2024b). With more information, the geographic distribution of suitable hosts could be overlayed onto the map of thermal suitability, to provide a more accurate representation of the PSHB range (i.e., from the fundamental niche to the realised niche). This has been done previously for PSHB in California using vegetation databases (Lynch et al., 2025; Umeda, 2017). A challenge here is that location data may not be available for many suitable host species. In particular, the ornamental plants favoured by PSHB may occur widely but are undocumented, planted in private properties (Umeda, 2017).

### Uncertainty of parameters estimates

Good model predictions rely on good parameter estimates. We parameterised our model using life history data for the *E. fornicatus* species complex gathered from multiple studies (Dodge, 2019; Gadd, 1941; Umeda and Paine, 2019; Walgama and Zalucki, 2007). Data for PSHB specifically was limited, so we had to assume similar thermal performance among the cryptic *E. fornicatus* species. The development rates estimated from Walgama and Zalucki, (2007), and Umeda and Paine, (2019), are remarkably consistent with one another. Walgama and Zalucki, (2007), measured nearly 100% survival of TSHB at the species’ optimal temperature, indicating a good husbandry setup that minimised confounding various. Laboratory studies rear the beetles on an artificial diet inoculated with the mutualistic fungus (Umeda and Paine, 2019; Walgama and Zalucki, 2007), and so our temperature performance curves (TPCs) inherently capture the performance of the *Euwallacea*-*Fusarium* symbiosis at different temperatures; a necessary consideration for mutualistic species (Hess et al., 2024).

Modelling temperature-dependent development needs to take into account the thermal conditions experienced by beetles within host sapwood. The microhabitat of wood-boring beetles can differ from the external air temperature by several degrees (Bolstad et al., 1997; Formby et al., 2018; Logan and Powell, 2001). Different locations within the same tree can vary greatly, however (e.g., the side of the trunk exposed to direct sunlight being much warmer than the shaded side; Bolstad et al., 1997; Logan and Powell, 2001; Powell, 1967). Additionally, the tree’s immediate surroundings and the species of tree itself (due to different thermal properties of wood, and the transpiration traits of the particular species) will influence temperature (Bolstad et al., 1997). We did not consider these factors as we were modelling over very large spatial scales, and instead wanted a generalised function for mean tree temperature. We fitted a model to actual sapwood measurements taken from sap flow loggers (Supp. Material). The sap logger dataset was limited, however (only for one location in Perth). A large set of measurements taken over a wide range of environments is needed to hone our microclimate model and to better understand the average response to mean climate conditions for a region. Unfortunately, accessible databases of internal tree temperatures appear to be lacking. Sap flow loggers would be a good source of data, and there are indeed extensive databases available from sap flow studies (e.g., Poyatos et al., 2016). However, the actual internal tree temperatures are not relevant for such studies, and as a consequence these data are often discarded.

The one life history parameter we were unable to estimate from the literature was *µ* – the loss of pre-oviposition adults during dispersal. This is very difficult to measure empirically. We can consider this value to be, in part, the product of the rate of dispersal and the mortality during dispersal. One study reported that females most frequently walk a short distance after emergence and establish a new gallery in the same stem, although some take to the wing (Gadd, 1949). Another laboratory study observed approximately 50% of females to fly away after emerging from a branch (Calnaido, 1965). The dispersal rate may be influenced by external factors such as time of day and rainfall (Calnaido, 1965; Liu et al., 2022). The probability of between-host dispersal is an uncertainty that needs to be quantified to better predict rates of transmission from infected hosts.

Mortality during dispersal is likely to be higher than the background mortality rate, since individuals are exposed to predators and other risks during this period. Additionally, there are many tree species in which PSHB will excavate a gallery but appear unable to establish a breeding colony, potentially due to the *Fusarium* fungus being unable to establish (DPIRD, 2024b; Mendel et al., 2021). These trees may act as ecological traps for PSHB, attracting females and thus preventing them from infesting a viable host. Mendel et al., (2021) found that only 20% of the host species attacked by invasive PSHB (in California and Israel) were suitable for reproduction; it is slightly higher (∼ 50%) for the current list of species attacked in Western Australia (DPIRD, 2024b). In the ambrosia beetle *Xyleborus saxeseni*, only 20% of observed galleries successfully produced offspring (Peer and Taborsky, 2007). The density in a landscape of non-reproductive hosts, as well as reproductive ones, will probably influence the rate of successful dispersal. Lynch et al., (2025) mapped the density of suitable PSHB hosts in California, calculated from the phylogenetic closeness of tree species to known susceptible species. Such an approach would not only provide a more accurate prediction of the species realised niche, it could also be used to map heterogeneity in dispersal/mortality rates arising from adults attacking non-reproductive hosts.

We honed our estimate of *µ* to within a reasonable window by running simulations under different *µ* values for invaded locations around the world (Figure 4). We found that for PSHB populations to be maintained at these sites, *µ* would need to be ≤0.4. This, of course, assumes the other aspects of the population model are reasonable for these locations. Small increases in *µ* made a large difference to population growth (Figure 4), which has interesting implications for PSHB management: treatments targeting dispersing females (through trapping, or ecological traps, for example) could have a large effect on population viability. A greater understanding of the drivers of dispersal will be valuable not only for improving our population model, but also for designing management strategies.

### Future directions

Our single-host stage-structured population model provides a foundation upon which more complex models for PSHB can be built. For example, the structure can be expanded to encompass multiple host trees, or multiple host communities (represented by grid cells), with dispersal of individuals occurring between them (Coates et al., 2022). Such a model can be used to simulate PSHB range expansion across a heterogeneous landscape over time, which would be a useful forecasting tool for risk mapping and to prepare for invasion into new areas (Malchow et al., 2024; Merow et al., 2011). Additionally, our model could be combined with optimisation models to determine how monitoring and eradication efforts should be allocated across space and time to achieve maximum success (Giljohann et al., 2011). Applying spatially-explicit decision models to our predicted distributions (Hauser and McCarthy, 2009) would provide quantifiable units to some of the management approaches that we suggest here – e.g., what proportion of containment efforts should be focused in high-suitability regions. Economic models also benefit from incorporating process models such as ours in their calculations. For example, an economic model estimating the cost of PSHB outbreaks and treatments in Western Australia assumed all areas are equally susceptible to PSHB (Cook and Broughton, 2023). These estimates – and in turn management decisions – could be refined by focusing on regions most at risk of PSHB invasion.

### Final remarks

PSHB invasions are an ongoing issue. The known range of this species has expanded with increased monitoring efforts into new parts of South Africa (Townsend et al., 2025, 2024) and from Israel into Palestine (Salman et al., 2019). The Quarantine Area in Perth, Australia, was extended in 2024, reflecting the difficulty in containing new outbreaks (DPIRD, 2024a). Furthermore, a new population of PSHB has very recently been detected in Spain (Goldarazena et al., 2025). Swift management decisions are needed to respond to the continuing spread of this species. Predictive models – especially those that can readily be applied to novel areas – are valuable tools to assist with making informed decision. The potential PSHB distribution generated by our SDM highlights key areas of concern, both within and outside the known range. By adopting a mechanistic framework to our approach, this model also predicts invasion dynamics of this species in greater detail. This includes estimating population growth rates, spread rates, seasonal dynamics and life history parameters. Our modelling framework can be expanded in the future to address specific management questions, such as how to deploy control strategies through space and time to achieve maximum success at eradication.

## Acknowledgements

We give our thanks to: David Owens and Paul Kendra for providing their PSHB mark-release-recapture data (Owens et al., 2019), which we used to estimate diffusion; Erik Veneklaas, Paul Drake, Wolfgang Lewandrowski and Emily Tudor for providing sap flow datasets which we used to predict tree temperature ; Xingyu (Mia) Li for providing PSHB distribution rasters generated by SDMs (Li et al., 2024) which we reproduced in this paper; and Nick Golding for assistance with the greta package for Bayesian inference.

## Data accessibility

Model code and associated datasets used to run these simulations can be found at GitHub (https://github.com/PopBiolGen/PSHB_Dynamics/releases/tag/PSHB_PopDyn).

The authors declare that they have no conflicts of interest.

