## Supplementary Material for "Boring beetles and super models: mapping the distribution of a new invader"

15/05/2025

The data used in the following material, and the scripts for reproducing it, can be found in the GitHub repository (https://github.com/PopBiolGen/PSHB_Dynamics), alongside the complete scripts for the PSHB model.

### PSHB Thermal Performance Curves (TPCs)

Three of our PSHB life-history parameters are temperature-dependent: the daily probabilities of juvenile transition ($\alpha_J$), pre-oviposition transition ($\alpha_P$) and juvenile survival ($\phi_J$). To generate functions for temperature-dependent transition and survival, we fit temperature performance curves (TPCs) to empirical PSHB data. TPCs describe the relationship between temperature, $T$, and performance, $P$ (in this instance, performance is the probability of transition or survival). Here, we use a TPC function as described in Phillips et al. (2014):

$$P(T) = \begin{cases} P_{max} e^{-\frac{-(T-T_{opt})^2}{2c^2}} & \text{for } T < T_{opt} \\ P_{max} e^{-\frac{-(T-T_{opt})^2}{2(cd)^2}} & \text{for } T \geq T_{opt} \end{cases} \quad \text{(Supp. Eq. 1)}$$

where $P_{max}$ is the maximum performance value and $T_{opt}$ is the optimum temperature (*i.e.*, the temperature at $P_{max}$). Performance increases with temperature up to $T_{opt}$, according to $c$. At temperatures above $T_{opt}$, performance decreases, as given by $d$.

TPCs for $\alpha_J$, $\alpha_P$ and $\phi_J$ were fit to empirical data, and values for $P_{max}$, $T_{opt}$, $c$ and $d$ were estimated (described below). When running the population model, the daily tree temperature was input as $T$ in Supp. Equation 1, to calculate the values of $\alpha_J$, $\alpha_P$ and $\phi_J$ to use for that time-step.

#### $\alpha_J$ TPC

The daily probability of juvenile PSHB transitioning to the preoviposition stage is given by $\alpha_J$. We estimated $\alpha_J$ using data from Umeda and Paine (2019), who monitored PSHB development (from egg to the pre-oviposition stage) under temperatures ranging 18 – 32 °C. Development rates are summarised in Supplementary Table 1. We also used data from Walgama and Zalucki (2007), who recorded development times for the tea shot hole borer (TSHB, within the *E. wallacea* species complex) at 15 – 32° C. Development times for eggs, larvae and pupae were obtained from Table 1, Figure 1, and Table 2, respectively (Supp. Table 1). For each temperature, we summed development times to calculate the total duration of the juvenile stage, $\delta$, and converted this to a daily transition rate: $\alpha_J = \delta^{-1}$ (Supp. Table 1). Transition rates from the two studies are plotted in Supp. Figure 1.

**Supplementary Table 1** The development times of PSHB eggs, larvae and pupae at different temperatures, from Walgama and Zalucki (2007), used to estimate daily juvenile transition rates; and daily juvenile transition rates at different temperatures, from Umeda and Paine (2019).

| Temperature (°C) | Egg development time (days) | Larva development time (days) | Pupa development time (days) | Juvenile transition rate (Walgama & Zalucki) | Juvenile transition rate (Umeda & Paine) |
|---|---|---|---|---|---|
| 15 | no hatching | | no emergence | 0 | |
| 18 | 23.5 | 34.72 | 15 | 0.014 | 0.013 |
| 20 | 18.5 | 25.06 | 14 | 0.017 | 0.02 |
| 22 | 13 | | 9 | | |
| 25 | 7.3 | 14.53 | 7.5 | 0.034 | 0.0258 |
| 28 | 5.5 | 7.14 | 6 | 0.054 | |
| 30 | 5 | | 4 | | 0.0421 |
| 32 | 10.9 | | 7 | | 0.0206 |

A TPC was fit to the empirical data using the maximum likelihood method (Supp. Figure 1). The mean parameter estimates for the TPC and their standard errors (solving the Hessian matrix and taking the square root) were:

- $P_{max} = 0.045(0.003)$
- $T_{opt} = 29.7(1.25)$
- $c = 6.9(1.07)$
- $d = 0.27(0.18)$

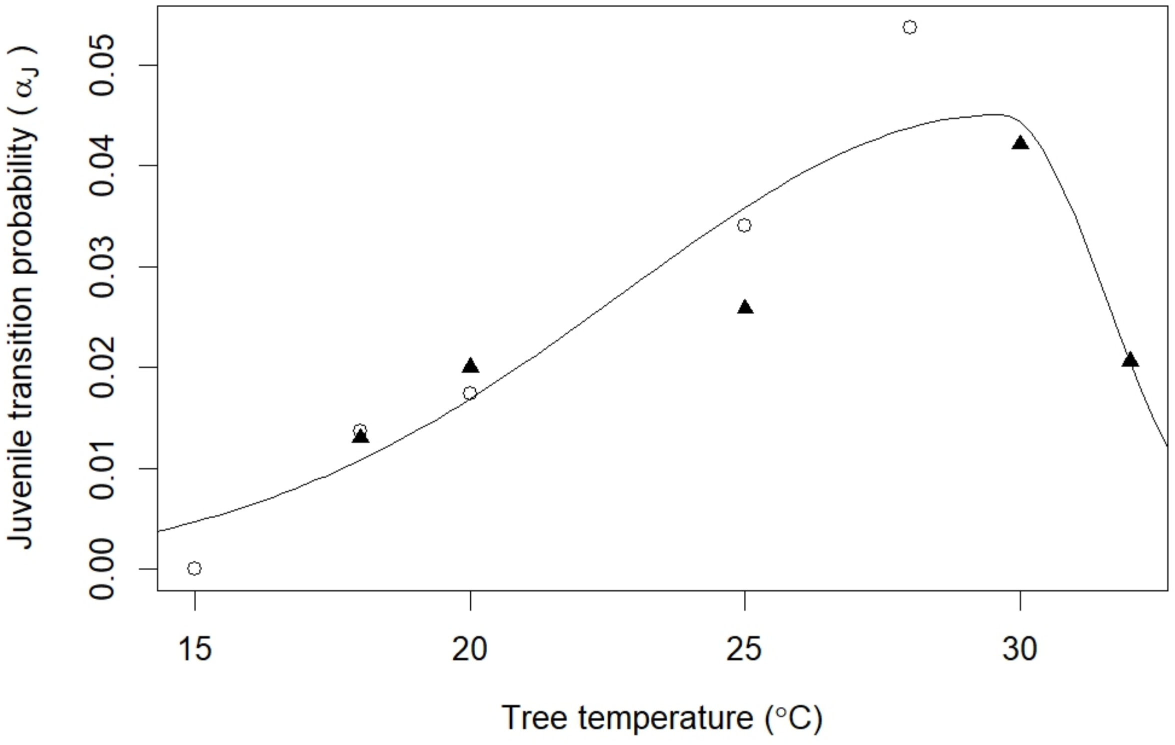


[]

**Supp. Fig. 1.** Juvenile transition probabilities at different temperatures, given by Umeda and Paine (2019) (triangles) and Walgama and Zalucki (2007) (circles), with the fitted TPC shown.

#### $\alpha_P$ TPC

The transition probability for the pre-ovipositing stage is given by $\alpha_P$. There is limited empirical data for this stage. In Walgama and Zalucki (2007), the pre-oviposition period for TSHB was estimated to be 136 degree-days, or 8 days at $T_{opt}$, and with a minimum temperature of 15°C. We therefore assigned $P_{max} = 1/8$. We assumed that the other TPC parameters ($T_{opt}$, $c$, $d$) were the same as for $\alpha_J$.

#### $\phi_J$ TPC

The daily survival probability of juveniles, $\phi_J$, is also supposed to be temperature-dependent. Walgama and Zalucki (2007) report the total egg and pupa mortalities at different temperatures. These data (extracted from Figure 2 of that study) are provided in Supp. Table 2. We used these values of total proportion mortality, along with the total duration of eggs and pupae at each temperature (Supp. Table 1), to calculate the daily probability of survival:

$$\phi = e^{-(\frac{-log(1-\lambda)}{\delta})} \qquad \text{(Supp. Eq. 2)}$$

where $\lambda$ is the total mortality of the stage, and $\delta$ is the total duration of the stage. Estimates of $\phi$ from the empirical data are given in Supp. Table 2, and plotted in Supp. Figure 2.

**Supplementary Table 2** Total proportion mortality of eggs and pupae at different temperatures (from Walgama and Zalucki (2007)), and the daily survival probabilities estimated from these data. Note that juveniles did not develop at 15 °C, so daily mortality could not be defined in Supp. Eq. 2.

| Temperature (°C) | Total egg mortality | Total pupa mortality | Daily egg survival | Daily pupa survival |
|---|---|---|---|---|
| 15 | 1 | 1 | | |
| 18 | 0.2 | 0.278 | 0.991 | 0.979 |
| 20 | 0.163 | 0.147 | 0.990 | 0.989 |
| 22 | 0.14 | 0.124 | 0.988 | 0.985 |
| 25 | 0.12 | 0.051 | 0.983 | 0.993 |
| 28 | 0.021 | 0.002 | 0.996 | 0.999 |
| 30 | 0.051 | 0.014 | 0.990 | 0.996 |
| 32 | 0.421 | 0.041 | 0.951 | 0.994 |

As above, we fitted a TPC to the empirical data (Supp. Fig. 2) and estimated the parameter values for the TPC function. The mean parameter estimates (and standard errors) were:

- $P_{max} = 0.99(0.004)$
- $T_{opt} = 29.5(2.36)$
- $c = 80(36)$
- $d = 0.15(0.19)$

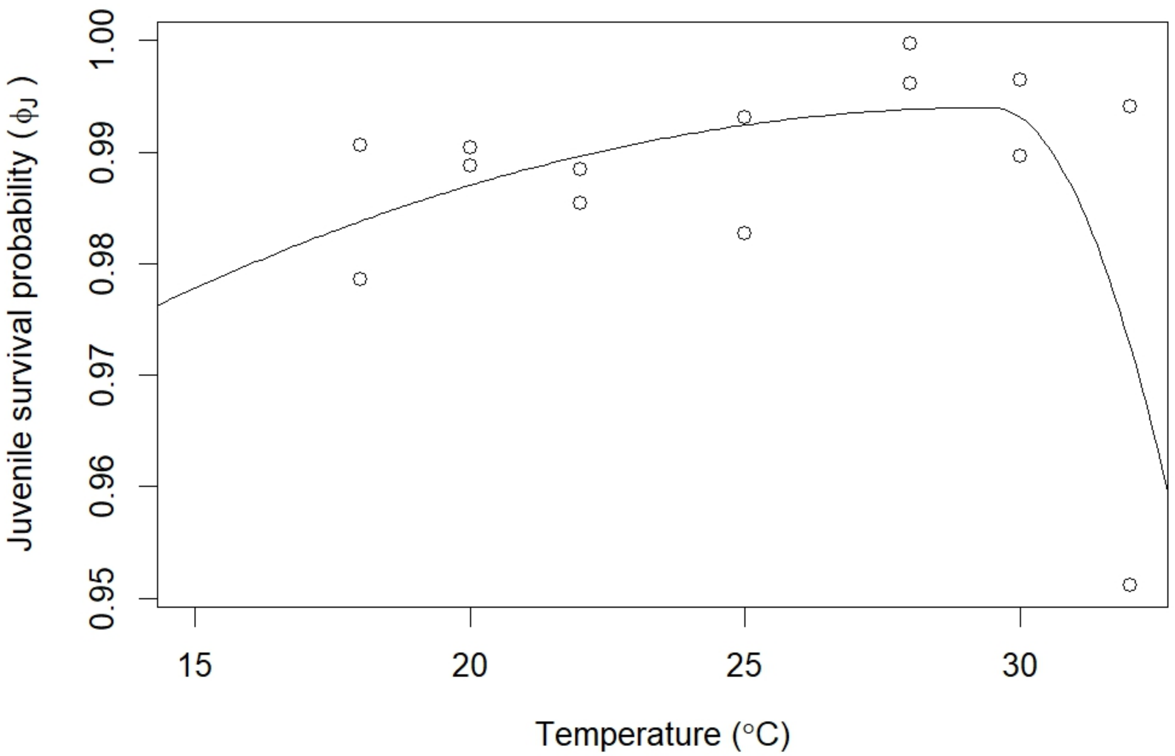


[]

**Supp. Fig. 2.** Juvenile survival probabilities at different temperatures given by Walgama and Zalucki (2007), with the fitted TPC function shown.

### Tree temperature predictions

Tree temperature, $T_{tree}$, was a weighted function of the maximum air temperature, $T_{max}$, and the soil temperature, $T_{soil}$:

$$T_{tree} = p_t T_{max} + (1 - p_t) T_{soil} \qquad \text{(Supp. Eq. 3)}$$

This assumes that the internal wood temperature does not exceed the maximum air temperature, nor get lower than the temperature of the soil. The relative weight of $T_{max}$ and $T_{soil}$ is given by $p_t$, which can take a value between 0 and 1. $p_t$ is a function of the relative humidity, $H$. Specifically, we assumed a linear relationship with humidity:

$$\text{logit}(p_t) = bH + c \qquad \text{(Supp. Eq. 4)}$$

To estimate values for $b$ and $c$, we fitted this model in a Bayesian framework using Hamiltonian Monte Carlo (HMC) Markov Chain, implemented in the greta package (Golding (2019)). Priors for $b$ and $c$ were set as normal distributions with mean = 0 and standard deviation = 3. The estimates (mean and standard errors) for the parameters in Supp Eq. 4 were calculated by samples from the posterior as:

- $b = 0.035(0.02)$

- $c = -0.48(0.7)$

When running the population model, tree temperatures were calculated each time-step according to Supp. Equations 3 and 4, using the above values of $b$ and $c$, and that day's $T_{tree}$, $T_{soil}$ and $H$ values (from the climate database).

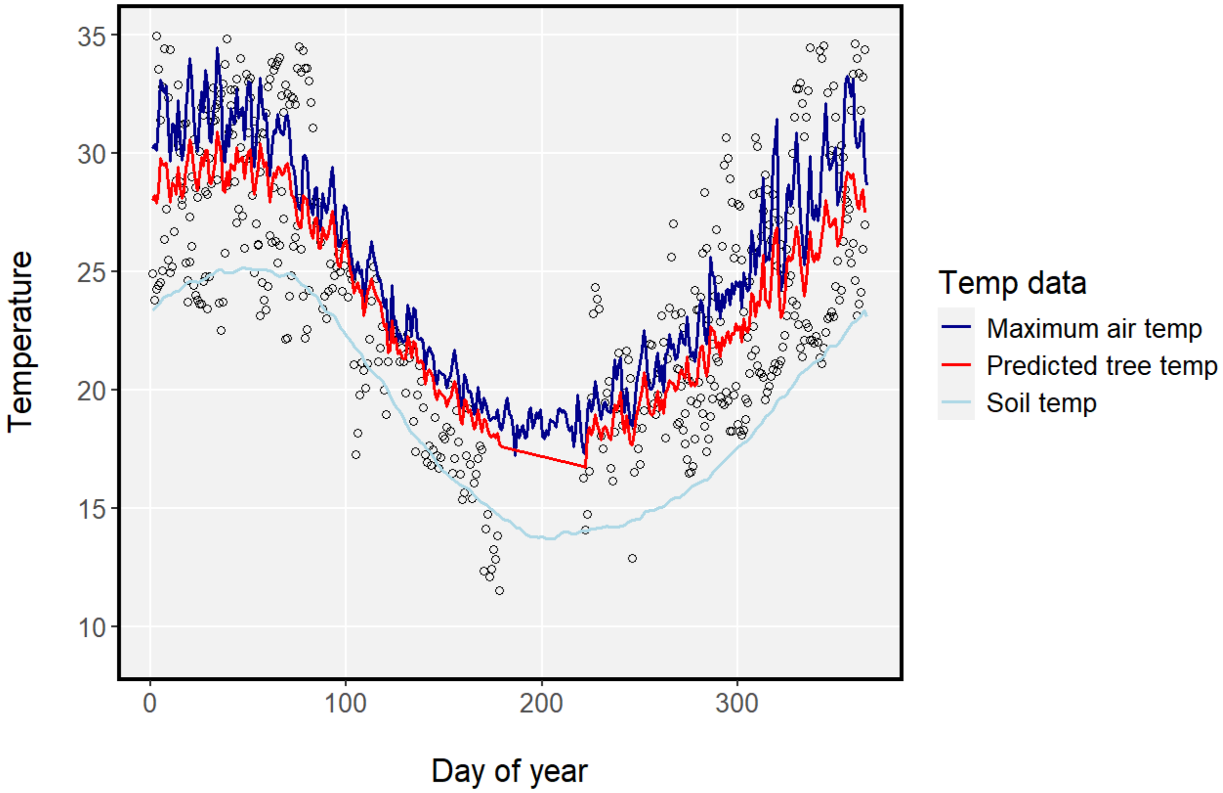


**Supp. Fig. 3.** Mean daily tree temperatures per day of year, as recorded by sap flow loggers (points), and as predicted by our model (red line), given the mean daily maximum air temperature for each day over the last decade (dark blue line) and mean soil temperature each day over the last decade (light blue).

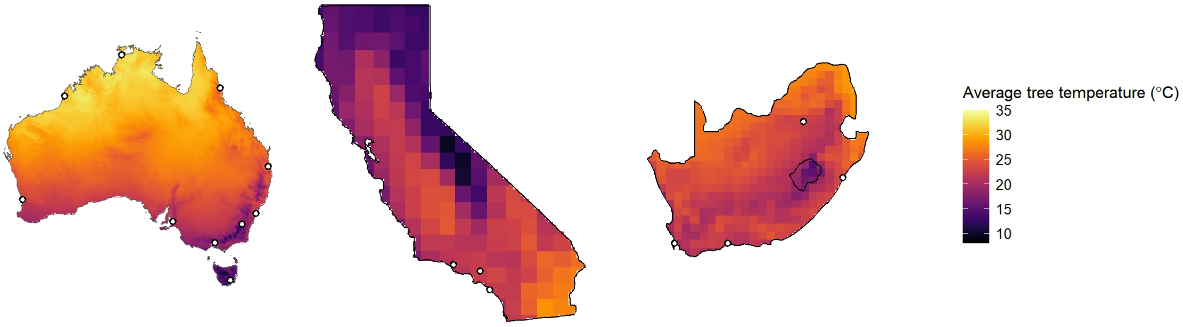


**Supp. Fig. 4.** Average tree temperature predicted for Australia, California and South Africa.

### Estimating diffusion coefficient

To estimate the diffusion coefficient, $D$, we generated a dispersal kernel by fitting a 2D Gaussian density function to mark-release-recapture data provided by Owens et al. (2019). The Euclidean distance, $d$, between the release point and each capture point was calculated. The number of individuals dispersing to a single point at distance $d$, given as $n(d)$, was expressed (in unsimplified form) as:

$$n(d) = \frac{n_0}{2\pi\sigma^2} e^{-\frac{d^2}{2\sigma^2}} \pi r^2 \qquad \text{(Supp. Eq. 5)}$$

Supp. Eq. 5 is the function for a 2D Gaussian distribution (Shigesada and Kawasaki (1997)), multiplied by $\pi r^2$ because we assumed each trap had a detection radius of $r$, whereby any individual dispersing to within $r$ of the point at $d$ will be counted. The parameter $n_0$ is the total number of dispersing individuals, which was set to $n_0 = 7007$ (from the mark-recapture data).

We used the Bayesian inference greta package to fit the model to the mark-recapture data, and to estimate values for $\sigma$ and $r$. We chose uninformative priors of $\sigma \sim \text{unif}(0, 300)$ and $r \sim \text{unif}(0, 100)$. The mean estimate from Bayesian inference was $\sigma = 25.7$, which we used to calculate the diffusion coefficient, as $D = \frac{\sigma^2}{2t}$ (Shigesada and Kawasaki (1997)).
